# Widespread of horizontal gene transfer events in eukaryotes

**DOI:** 10.1101/2022.07.26.501571

**Authors:** Kun Li, Fazhe Yan, Zhongqu Duan, David L. Adelson, Chaochun Wei

## Abstract

Horizontal gene transfer (HGT) is the transfer of genetic material between distantly related organisms. While most genes in prokaryotes can be horizontally transferred, HGT events in eukaryotes are considered as rare, particularly in mammals. Here we reported the identification of HGT regions (HGTs), which are genomic sequence fragments indicating the occurrence of HGT events, in human, mouse, cow, lizard, frog, zebrafish, fruit fly, nematode, Arabidopsis and yeast. By comparing the genomes of these 10 representative eukaryotes with 1,496 eukaryotic genomes, 16,098 bacteria and 11,695 viruses, we found between 10 and 243 non-redundant HGTs per species, and most of these HGTs were previously unknown. These HGTs have transformed their host genomes with various numbers of copies and have impacted hundreds, even thousands of genes. We listed several examples of HGTs and proposed some possible routes that HGT events occurred. Further analysis showed that the majority of the 1,496 eukaryotes with full length genome sequences also contain HGTs. Our findings reveal that HGT is widespread in eukaryotic genomes, and HGT is a ubiquitous driver of genome evolution for eukaryotes.

## Background

Horizontal gene transfer (HGT) is the transfer of genetic material between organisms that is not from parent to offspring, and it is a major driver of genome evolution in bacteria and archaea[1, 2]. On average, 81% of genes in bacteria were involved in HGT[3]. Recent evidence has shown that HGT events also exist in eukaryotes. For example, HGT regions (HGTs) have been reported from soil bacteria to the common ancestor of *Zygnematophyceae* and *embryophytes*, which increased its resistance to biotic and abiotic stresses during terrestrial adaptation[4]. HGT of a plant detoxification gene BtPMaT1, made whiteflies gain the ability to malonylate a common group of plant defense compounds[5]. Besides, 1,410 genes were acquired via 741 distinct transfers from non-metazoan donors in 218 insects and males lacking LOC105383139 which was transferred from bacteria courted females significantly less in diamondback moths[6]. Another remarkable example of HGT is a ∼1.5Mb fragment of *Wolbachia spp.* DNA integrated into the pill bug *Armadillidium vulgare* genome, resulting in the creation of a new W sex chromosome[7]. HGTs have been observed in genomes of five parasitic plants in the *Orobanchaceae* family[8], several unicellular pathogens[9] and blood-sucking parasites[10–12]. It has been proposed that only unicellular and early developmental stages of eukaryotes are vulnerable to HGT[13], while some argue that HGT events in eukaryotes may be limited to those derived from endosymbiotic organelles[14, 15]. Mechanisms for the transfer of DNA into eukaryotic genomes have been described for viral infection, transposons, conjugation between bacteria and eukaryotes, or from endosymbionts (not only plastids and mitochondria)[16]. Some behaviors, such as predation, and life-styles, such as parasitism, have been reported to promote DNA transfer in eukaryotes[17]. Therefore, while the prevalence of HGT may be rare in eukaryotes, compared to bacteria and archaea, it does occur. However, the scale and impact of HGT events in eukaryotes are unknown.

We presented here a fast identification method for HGTs in eukaryotes using both sequence composition bias and genome comparison. We evaluated the method by a simulated dataset and a dataset of HGT regions reported previously for whitefly. We applied this method to 10 representative organisms with high quality genomes which were at different positions in phylogenetic tree of eukaryotes. Many bacterial and viral genomes were also compared to reveal the potential media organisms or vectors for HGT in eukaryotes.

## Results

### A Fast HGT identification method and its evaluation

We created a fast identification pipeline for HGTs in eukaryotes by combining sequence composition filtering and sequence alignment (Figure S1; see Methods). In brief, we first identified genomic regions most different from the rest of the genome based on their k-mer frequencies and then we aligned these selected genomic regions to other genomes. Before sequence alignment, we divided all genomes into three different groups based on their taxonomic information, including self group (SG), closely related group (CRG) and distantly related group (DRG). The genomic regions were considered as candidate HGT regions if their sequence conservation levels were discordant to the phylogenetic tree of relevant species. Specifically, genomic fragments were determined as HGT sequences occurred in the common ancestor of organisms listed in SG if they had higher identity percentage with species in DRG than that in CRG. To further evaluate the accuracy of our HGT identification pipeline, we evaluated it with a simulated dataset and a real dataset (HGTs reported in whitefly).

We evaluated the pipeline with a genome containing simulated HGTs (see Methods). Since our HGT identification pipeline has two main steps (sequence composition-based filtering and sequence alignment), the evaluation was done for the two steps. While top 1% fragments were input to the pipeline, 20.6% correct results would be identified after sequence composition-based filtering and 17.77% correct results identified after sequence alignment (Figure S2A; Table S1). When the percentage of fragments input was up to 20%, 58.9% and 57.1% correct results were identified after two steps respectively and it can be seen that the precision of prediction was higher than 90% for all cases (Figure S2A; Table S1). This indicates that we may have underestimated the number of HGTs (low recall rate) but majority of the identified HGTs are reliable.

We also evaluated our pipeline on the 170 HGTs previously reported for whitefly [6] (see Methods). While only top 20% genome fragments were input to our pipeline, 71.8% of the 170 HGTs would be identified after sequence composition-based filtering, indicating that the first step is effective to keep candidate HGTs, and 32.4% of the 170 HGTs would be identified after sequence-alignment-step (Figure S2A), indicating that our nucleotide sequence comparison based HGT identification method is different with existing methods. Overall, 131 HGTs were identified by our pipeline, 47.3% of which (62 HGTs) were overlapped with reported HGTs while 52.7% of which (69 HGTs) were not reported (Figure S2B; Table S2). Among 69 HGTs not reported previously, 36 HGTs did not overlap with protein-coding sequences, which made them invisible to the previous reported HGT identification methods limited to protein coding regions. The remaining 33 newly found HGTs did overlap with protein-coding regions, but the coverage of these protein coding regions by HGTs was less than 25% of their corresponding total protein, which might be the reason why they were not identified by previous method (Figure S2B). For the 115 HGTs reported previously but missed by our method, 48 HGTs were left out by sequence composition-based filtering step and the remaining 67 HGTs were not identified by our pipeline. Among them, 35 had more similar sequences with genomes in CRG than that in DRG while the other 32 had similarity lower than 50% with genomes in DRG (Figure S2B).

The comparison with previously reported HGTs show that our results are reliable and our method could identify some novel HGT events because it broke the restriction of identifying HGT regions involving protein coding regions only.

### Widespread of HGT events among eukaryotes

We applied our HGT identification method to identify HGTs in 10 representative organisms with high quality genomes, including 1 primate, 2 mammals, 3 non-mammalian vertebrates, 2 invertebrates, 1 plant and 1 fungus. We identified between 10 and 243 non-redundant cross-kingdom eukaryotic HGTs for these 10 representative organisms (Figure 1A; Table S3). The number of HGTs found in nematode, an invertebrate, was the smallest, while the number for Arabidopsis was the largest (Figure 1A). HGT events occurred to the common ancestors of organisms contained in self groups (SGs) of each representative organism. The HGTs of nematode mostly occurred at species level, while the HGTs of fruit fly and yeast mostly occurred at order level; human, mouse and Arabidopsis had more HGTs at class level, while cow, lizard, frog and zebrafish had more HGTs at phylum level (Figure 1A). The cross-kingdom species having the most similar sequence with the target organism was identified as the HGT related organisms. Overall, the HGT related organisms of these 10 representative species were distributed in 6 different kingdoms (bacteria: 196/745 [26.3%], metazoan: 168/745 [22.6%], protozoa: 154/745 [20.7%], plants: 153/745 [20.5%], fungi: 66/745 [8.9%], viruses: 8/745 [1.1%]) (Figure 1B; Table S4). Bacteria, metazoan, plants and protozoa occupied the highest proportion of HGT related organisms in mammals (human, mouse, cow), non-mammalian metazoan (lizard, frog, zebrafish, fruit fly and nematode) and non-metazoan eukaryotes (yeast and Arabidopsis) respectively (Figure 1B). Especially, the *Actinomycetes* of bacteria, the *Insecta* of metazoan, the *Magnoliatae* of plants, the *Aconoidasida* of protozoa and the *Sordariomycetes* of fungi contained prevalent HGT related organisms (Figure S3). Besides, most of the identified HGTs in the 10 representative organisms were previously unknown compared with reported HGTs[11, 18–25] (Figure 1C).

**Figure 1.**
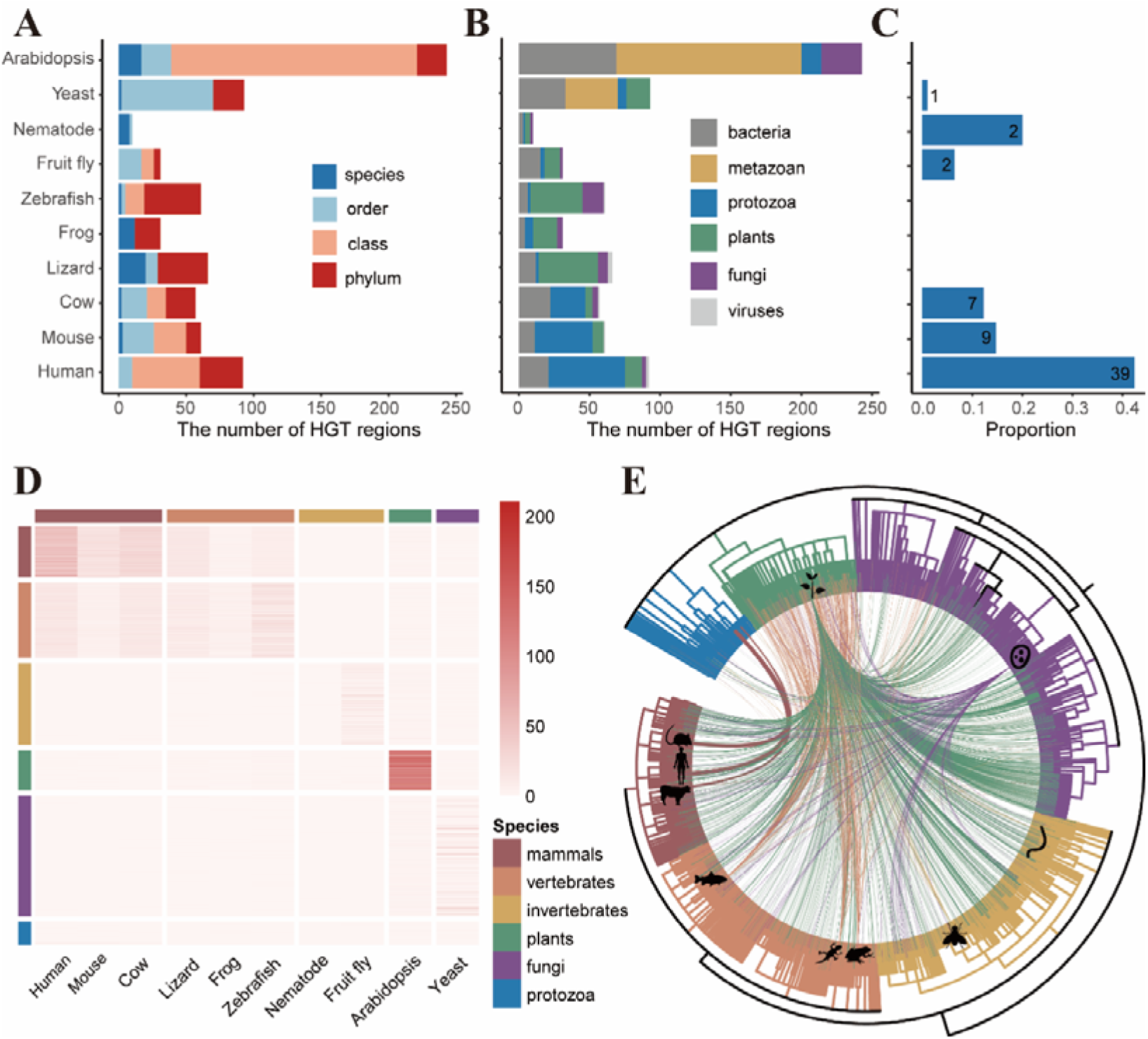
745 HGTs identified in 10 representative organisms. (A) The number of HGTs occurred to the common ancestors of organisms contained in different self groups (SGs) for 10 representative organisms. Organisms of a certain taxonomy level, such as species, order, class and phylum, can be considered as a self group. (B) The number of HGTs with different HGT donor/acceptor organisms in 10 representative organisms. (C) The proportion of HGTs reported previously[11, 18-25]. (D) The HGT-appearance numbers between 10 representative organisms (X axis) and 1,496 eukaryotes (Y axis) are represented by the grid colors in the heat map. (E) HGTs between eukaryotes were shown by the lines connecting related species, and the thickness of the line represented the HGT-appearance number of the related species.

To determine the frequency of HGT in eukaryotes, we calculated an HGT-appearance number N_AB_ for a representative organism A and another eukaryotic organism B, which was defined by the frequency with which organism B appeared in the phylogenetic trees of non-redundant HGTs of representative organism A. For instance, among the 92 non-redundant HGT trees for Homo sapiens, Pan troglodytes was found in 89 of them, therefore the HGT-appearance number N_AB_ between Homo sapiens and Pan troglodytes was 89. The distribution of HGT-appearance numbers N_AB_ between the 10 representative organisms and 1496 eukaryotes was shown in Figure 1D and Table S5. The HGT-appearance numbers of organisms having the same taxonomic classification were higher (Figure 1D), indicating that most HGTs identified by our pipeline were transferred before the divergence of model organisms and their sibling lineages. This also implies that these HGTs may have important functions since they have survived during the evolution[1]. By using this metric, we determined that 90.2% of eukaryotes (1349 of 1496) hosted HGTs (Figure 1D, 1E), revealing widespread of HGT events across eukaryotes.

### Duplications of HGTs and their impact on their host genomes

We compared the non-redundant eukaryotic HGTs we identified with their host genomes to further elucidate the impact of HGTs on their host genomes. Overall, 77.2% of HGTs (575 of 745) have a single copy in their host genomes, and the rest (22.8%, 170 of 745) have multiple copies. Especially, 6 HGTs have more than 100 copies (Figure 2A; Table S3). In particular, HGT region “NC_037357.1:83623202-83624198” related to BovB in the cow genome had 36,868 copies (with a total length of 36.3 Mbps), which was consistent with a previous study that BovB were present as many copies[10]. In newly identified HGTs, zebrafish HGT region “NW_018395302.1-126431-127344” had 126 copies and occupied 18 Kbps.

**Figure 2.**
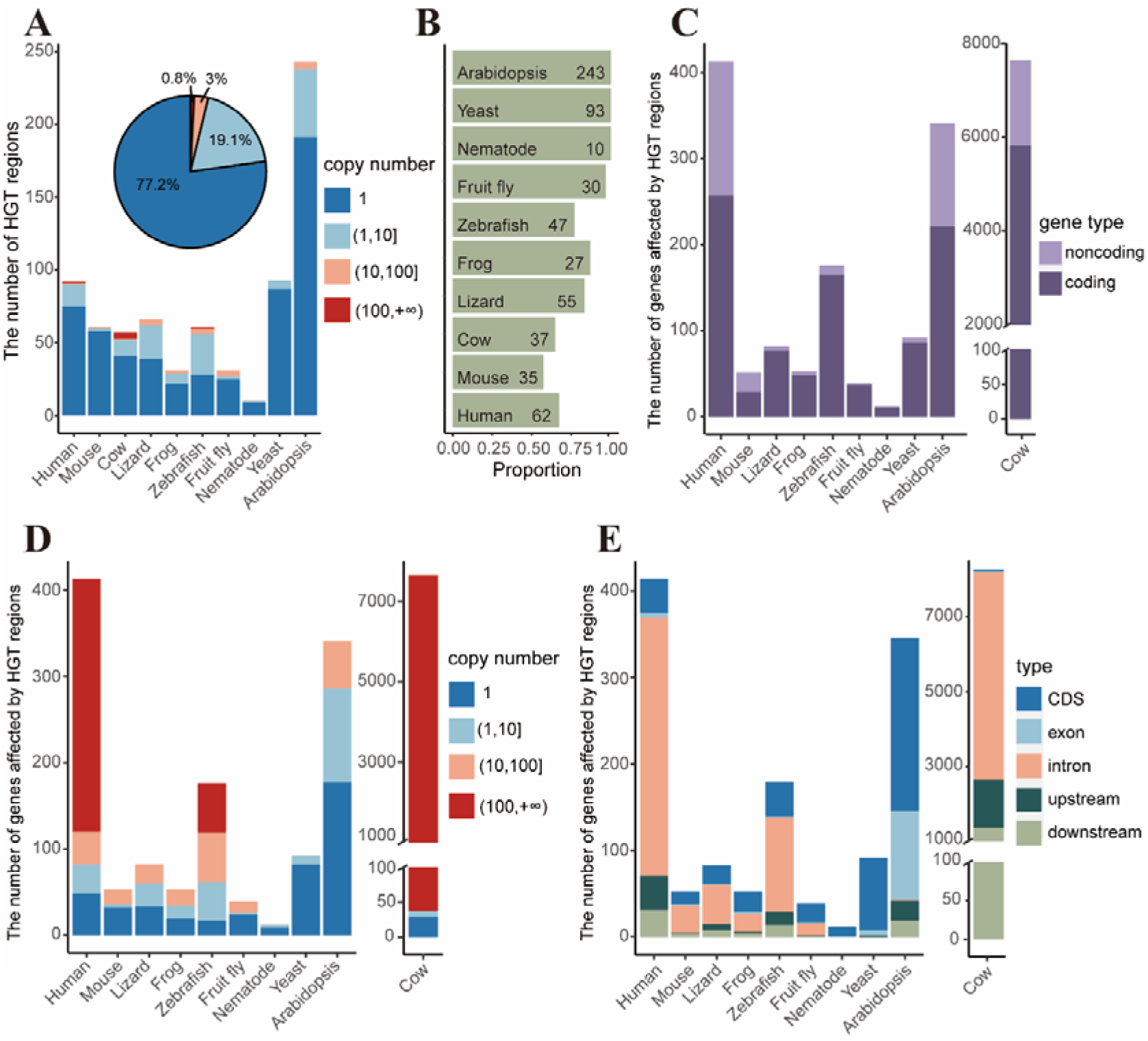
Duplications of HGTs and their impact on 10 representative organisms. (A) The composition of HGTs with different copy numbers. Although the number of unique HGTs were bigger, some HGT regions can have a huge number of duplications. (B) The proportion of HGTs overlapped with genes. Majority of the HGTs overlap with protein coding gene regions in most organisms except the mouse. (C) The number of genes affected by HGT regions, not only in protein coding genes, but also in noncoding genes. (D) The number of genes affected by HGT regions with different copy numbers. Human, cow and zebrafish genomes have hundreds or even thousands of genes impacted by HGT duplications. (E) The number of genic elements affected by HGT regions. Most HGT regions were in introns in vertebrates and fruit fly, while in worm, fungi and plants, most of the HGT regions were in CDS regions.

These HGT copies had affected many genes as well. In total, 93% of HGTs (693 of 745) had influence on 8,910 gene regions and 75.8% of them were overlapping with protein coding genes (Figure 2B and 2C). Especially, almost all HGTs impacted genes in Arabidopsis, yeast, nematode and fruit fly. There were 16 HGTs, each of which impacted more than 10 genes in their host genome (Table S3). For example, the zebrafish HGT region mentioned above and its copies overlapped (at least 1bp) with 52 protein-coding genes (Figure 2D), which was a huge impact on the zebrafish genome functions. HGTs with similar (but different degree of) impact on genome can also be found for most of the 10 representative organisms. Gene ontology (GO) analysis of genes affected by HGTs in different organisms shows that some were associated with ion channel (cow), synaptic membrane (human), actin cytoskeleton (mouse), thioester biosynthetic process (fruit fly), organic hydroxy compound metabolic process (yeast), biosynthetic process (Arabidopsis), etc. (Figure S3; Table S6). The genes impacted by HGTs in the remaining 4 of the 10 representative species have no enriched functions.

The genes mentioned above were affected by HGTs in different genetic elements. As a whole, only 5.2% (495/8.905) of genes overlapped by HGTs were affected in their CDS regions, with the rest were affected in the intron, upstream, downstream and exon regions (Figure 2E). This was because HGTs with a large copy numbers were mostly affected the intron regions of the genes (Figure S5), which might conform a rule for the occurrence of HGT, ‘first do no harm’[1]. In details, 75.3% (354/470) of genes affected by single-copy HGTs were affected in their CDS regions, while this proportion dropped to 0.2% (141/8435) for multi-copy HGTs and 70.9% (5984/8435) of genes affected by multi-copy HGTs were affected in their intron regions. Besides, the distribution of gene elements affected by copies of HGTs was different in different organisms (Figure 2E). Most copies of HGTs overlapped with introns in vertebrates, while in invertebrates, fungi and plants, most of the HGT regions overlapped with CDS regions.

### Repetitive sequence composition of HGTs

We compared the non-redundant HGTs detected in 10 representative organisms with the repetitive sequences annotated in their reference genomes. Between 0∼77% of their HGTs overlapped with interspersed repeats (excluding simple repeats) (Figure 3A; Table S7), revealing significant species and repeat-specificity. The types of repeats overlapping with HGTs showed significant correlation with overall genomic repeat composition. Long interspersed nuclear elements (LINEs) retrotransposons were common in HGTs detected in mammals, frog and lizard, while long terminal repeat (LTR) retrotransposons were common in HGTs detected in fruit fly, yeast and Arabidopsis, consistent with their frequencies in their host genomes (Figure 3B and S6). In comparison, the distribution of LTR in zebrafish HGTs was not consistent with their distribution in the host genomes, as seen the LTR in frog HGTs and the satellite in cow HGTs (Figure 3B and S6). In the zebrafish genome, LTR appeared in as many as 37 non-redundant HGTs (60.7%), while that repeat only accounted for 4.2% of repeats in the genome.

**Figure 3.**
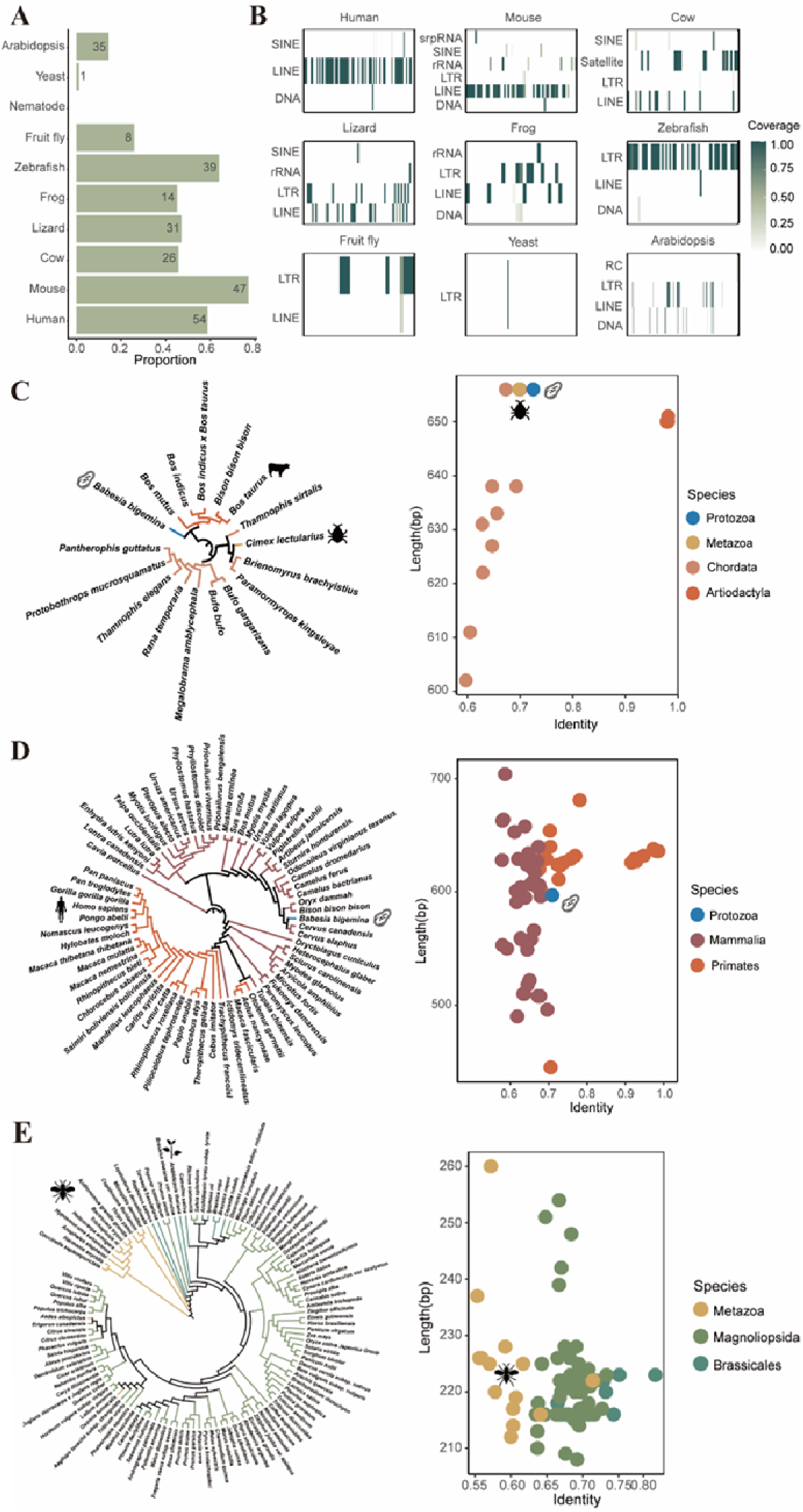
HGTs overlapping repetitive regions. (A) The proportion of HGTs overlapping with repeat sequences. Between 0∼77% HGTs overlapped with interspersed repeats. (B) The distribution of repeat sequence types in HGTs. LINE retrotransposons were common in HGTs detected in mammals, frog and lizard, while LTR retrotransposons were common in zebrafish, fruit fly, yeast and Arabidopsis. Phylogenic trees and length-vs-identity plots of HGTs. The trees on left side represent the evolutionary relationship of species related to this HGT region, and the plots on right present sequence similarity between the homologous sequences from the model organism and the related species. (C) Cow HGT region “NC_037344.1: 42095551-42096199” overlapped with BovB retrotransposons; (D) Human HGT region “NC_000004.12:121865648-121866285” overlapped with L1 retrotransposons; and (E) Arabidopsis HGT region “NC_003076.8:12073716-12073960” overlapped with Gypsy LTR retrotransposons.

BovB and L1 retrotransposons are the two most abundant transposable elements (TEs) in eukaryotes and replicate via an RNA intermediate[26]. The horizontal transfer of BovB is known to be widespread in animals[11] and horizontal transfer of L1 has been shown in plants, animals and several fungi[18]. In total, 4 of our non-redundant HGT events overlapped with BovB retrotransposons in cow (Table S8), supporting previous results for horizontal transfer of BovB[11, 18]. Furthermore, 119 L1 horizontal transfer events were identified in human, mouse, cow, lizard, frog and zebrafish (Table S8), providing more evidence that L1 elements are horizontally transferred[18].

All HGTs that overlapped with BovB retrotransposon were associated with the intracellular pathogen *Babesia bigemina* while one of them was also associated with the blood-sucking parasite *Cimex lectularius* (bed bug) (Table S8), both of which were the reported possible intermediary species[12]. *Babesia bigemina* is an Apicomplexan parasite that infects red blood cells [27], which infects livestock worldwide, including wild and domestic vertebrate animals. *Cimex lectularius* is known to feed on animal blood and can host over 40 zoonotic pathogens[28], thus transmitting many infectious diseases[29]. Figure 3C showed the tree of cow HGT region “NC_037344.1: 42095551-42096199” and its homologs. In addition to the two candidate vector species, this HGT tree also included 5 bovid mammals and 10 non-mammalian vertebrates (3 fishes, 3 amphibians, 4 reptiles) (Figure 3C; Table S9), which were clearly clustered in distinct branches. Like in other parasites[10], it appears that *Cimex lectularius* and *Babesia bigemina* transfer DNA between the hosts it feeds on.

A considerable number of genes of intracellular pathogens have been acquired through HGT, including Apicomplexans[30, 31]. Our analysis identified 76.5% (91/119) HGTs overlapped with L1 retrotransposons were associated with Apicomplexan intracellular pathogens, including *Babesia bigemina* and *Plasmodium vivax* (Table S8). For example, the tree of human HGT region “NC_000004.12:121865648-121866285” (Figure 3D; Table S9) had shown the transfer between *Babesia bigemina* and mammals, including 26 primates, 11 Artiodactyls, 11 Carnivores, 9 Rodents, 8 Chiropterans, 1 Eulipotyphl, 1 Lagomorph and 1 tree shrew.

Gypsy LTR retrotransposons are widely distributed among eukaryotes and have been found in plants, fungi and vertebrates[32]. The HGT of Gypsy is well known in Drosophila[33] and plants[34], and has been found between fungi and non-seed plants[35]. In our results, 73 HGTs overlapped with Gypsy were identified in zebrafish, Arabidopsis, lizard, frog, fruit fly and yeast, which further supported previous reports. The Arabidopsis HGT region “NC_003076.8:12073716-12073960” showed the horizontal transmission of Gypsy LTR retrotransposons took place among 105 magnoliopsida plants and 14 insects (Figure 3E; Table S9). Some HGTs overlapping with repeat sequences of other 6 representative organisms were in Figure S7.

### HGTs overlapping with non-repeating sequences

The above examples were mainly HGTs related to transposons and occurred in protozoa and eukaryotes. There were also 490 HGTs not overlapped with repetitive sequences (Figure 3A; Table S7), and 433 HGTs (88.4%) of them overlapped with 530 protein coding genes (Table S10) while 90.9% (482/530) of protein coding genes affected by 433 HGTs were affected in CDS regions (Figure 4A; Table S10). Gene ontology (GO) analysis of protein coding genes affected by HGTs in different organisms showed that some were associated with aldo−keto reductase activity (human), lipid oxidation (mouse), organic hydroxy compound metabolic process (yeast), organophosphate biosynthetic process (Arabidopsis), etc. (Figure S8; Table S11). We did not find enriched functions for the protein coding genes impacted by HGTs in the remaining 6 of the 10 representative species.

**Figure 4.**
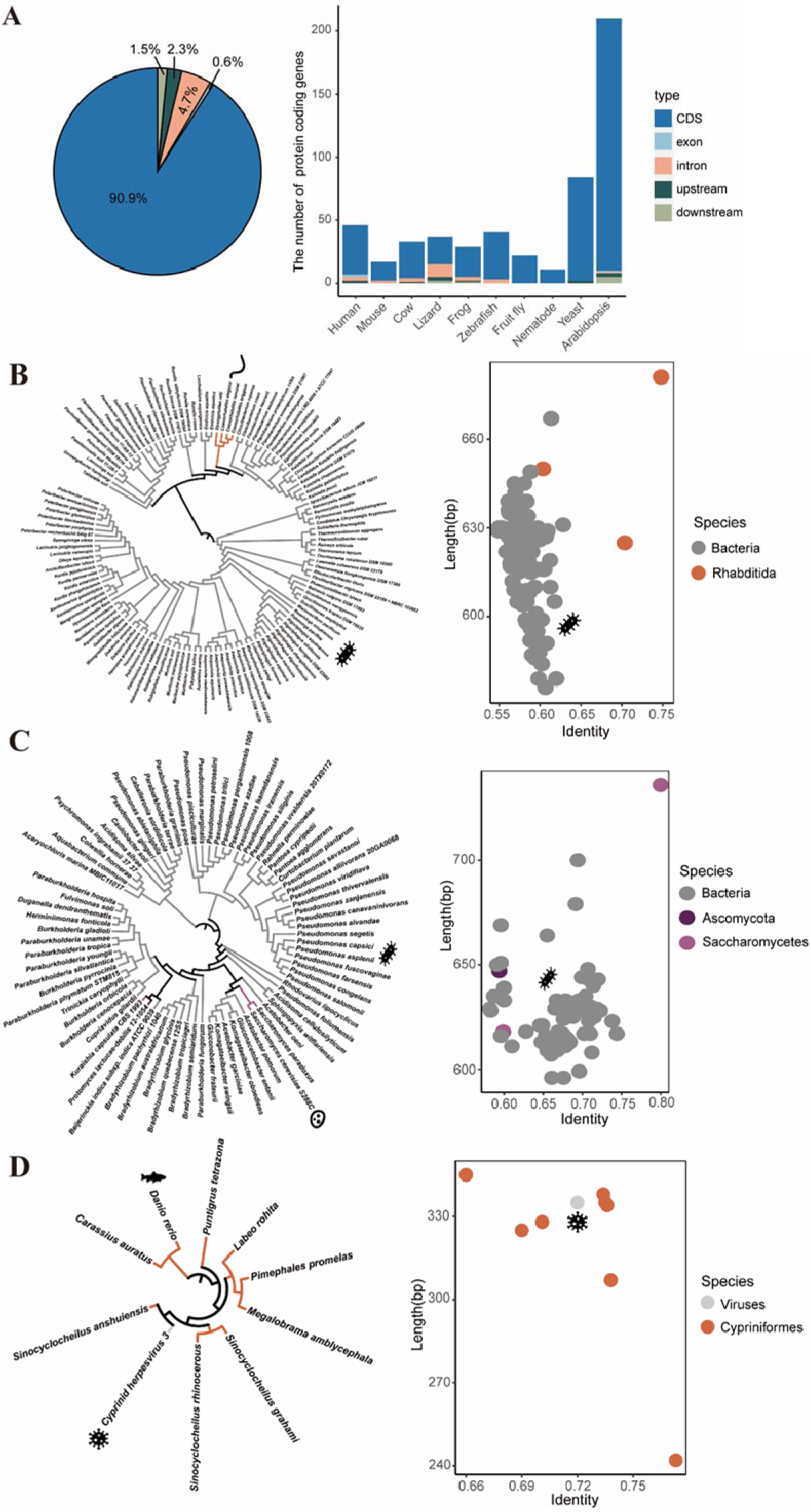
HGTs not overlapping repetitive regions. (A) The proportion of regions of protein coding genes affected by HGTs not overlapping with repeat sequences. Most protein coding genes affected by HGT events not overlapping with repeat sequences were affected on CDS regions. Phylogenic trees and length-vs-identity plots of HGT regions not overlapping with repetitive regions. The trees on left side represent the evolutionary relationship of species linked by this HGT region, and the plots present sequence similarity between the homologous sequences from the model organism and the related species. (B) Nematode HGT region “NC_003280.10:13635039-13635734” overlapped with gene CELE_C31C9.1 encoding a gut apical protein located in membrane raft; (C) Yeast HGT region “NC_001144.5:21121-21842” overlapped with gene YLL060C encoding Glutathione transferase GTT2; and (D) Zebrafish HGT region “NC_007123.7:45453300-45453643” overlapped with gene LOC562542, whose function is not clear.

In order to better understand HGT events, we also listed several HGTs not overlapped with repetitive sequences. These HGTs were associated with bacteria and viruses. The HGT region “NC_003280.10:13635039-13635734” of nematode suggested the gene transfer between rhabditida animals and bacteria (Figure 4A; Table S9) and it overlapped with the CDS of gene CELE_C31C9.1, which encodes a gut apical protein located in membrane raft[36]. The yeast HGT region “NC_001144.5:21121-21842” was overlapped with the CDS of gene YLL060C, having homologous sequences with ascomycota fungi and pseudomonadota bacteria (Figure 4B; Table S9). Glutathione transferase GTT2 encoded by gene YLL060C is a novel atypical type of cytosolic Glutathione-S-transferases (GSTs), which are detoxification enzymes that catalyse the conjugation of electrophilic substrates to glutathione[37]. The HGT region “NC_007123.7:45453300-45453643” of zebrafish suggested the gene transfer between cyprinoid fishes and viruses (Figure 4C; Table S9) and the HGT was overlapped with the CDS of gene LOC562542, whose function is not clear. More trees of HGTs can be founded in Table S10. Some HGTs not overlapping with repeat sequences but overlapping with CDS regions of protein coding genes of other 7 representative organisms were in Figure S9.

## Discussion

In this study, we created a fast identification pipeline for HGTs in eukaryotes by combining sequence composition bias and sequence alignment and applied this pipeline to 10 representative organisms. 745 HGTs were identified in the 10 model organisms and 90.2% of eukaryotes with whole genome sequences were involved in HGT, which suggests HGT events are widespread in eukaryotes. Compared to HGT in prokaryotes, the number of non-redundant eukaryotic HGTs (10∼243 regions) detected in these model organisms was very small. We found many HGT regions by comparing a small part of the genome sequences that were significantly different from their reference genomes. Besides, all HGTs we found were cross-kingdom, which means HGT only occurred between species in the same kingdom is not in our consideration. It is conceivable that the number of HGT regions is much larger than this.

In addition, most HGT regions identified by our pipeline were transferred before the divergence of model organisms and their sibling lineages, which implied that these HGTs may have important functions as they have persisted. Overall, 22.8% of HGTs had multiple copies in their host genomes and these HGTs and their copies overlapped with 8,910 genes. Both of them indicate that HGT events affect not only genome size but also genome functions. However, only 5.2% of genes were effected by HGTs in their CDS regions with the rest affected in the intron, upstream, downstream and exon regions. This result also indicates that it is reasonable to identify HGTs using genomic sequences, which can find some HGTs in non-coding regions compared to methods focusing on protein regions only.

Between 0∼77% of HGTs overlapped with interspersed repeats, in which BovB, L1 and Gypsy retrotransposons had the highest frequency. By analyzing the phylogenic tree of those HGTs, we propose that blood-sucking parasites (*Cimex lectularius*) and intracellular pathogens (*Babesia bigemina*) are involved in BovB transfer between mammals and other vertebrates, while Apicomplexans (*Babesia bigemina* and *Plasmodium vivax*) are the hotspot of L1 transfer in eukaryotes, and Gypsy can be transferred between insects and plants. However, the routes for DNA transfer for the majority of HGT events in this report are still unclear. With the progress of sequencing technology, especially third generation sequencing technologies, high quality whole genome sequences could be obtained for several HGT related species distributed across the tree of life, and this would provide a good opportunity to determine the route and direction of HGT.

In conclusion, comparison of 10 representative eukaryote genomes against other organisms with whole genome sequences available shows that HGT is widespread in eukaryotes. HGTs and their copies affect not only genome size but also genome functions. We list several HGTs overlapping with retrotransposons or having functions and propose some possible routes for these HGT events. Although we have only analyzed HGT events for 10 representative species, it is enough to see that HGT events vary greatly between species. There is still much work to be done to better understand HGT.

## Methods

In bacterial genomes, HGT regions are also called genomic islands (GIs) and can be detected using two distinct bioinformatics approaches, based on sequence composition or sequence alignment[38]. In general, the sequence composition of GIs is significantly different from that of the recipient genome. Composition-based methods identify GIs within genome sequences by calculating the k-mer frequencies of a fragment and comparing that frequency distribution with that obtained from the whole genome. Sequence alignment approaches are based on the premise that DNA sequence based phylogenetic tree topology of GIs will be discordant with respect to known species relationships, where sequences that are absent in several closely related organisms appear in more distant species. These two methods can be adapted to the identification of HGTs in eukaryotes but not without challenges. Due to the large sizes and the high heterogeneity of eukaryotic genomes, composition-based approaches may produce a number of false-positive predictions while sequence alignment methods are computationally expensive and time-consuming when hundreds of reference genomes must be aligned. In this study, we identified HGTs between eukaryotes and other organisms by combining these two approaches to reduce both the false-positive rate and computational cost.

### Data collection

Three datasets were downloaded from NCBI Refseq database[39] (ftp://ftp.ncbi.nlm.nih.gov/genomes/refseq). The first dataset contained the reference genome sequences of 10 representative organisms consisting of 3 mammals, 3 non-mammalian vertebrates, 2 invertebrates, 1 fungus and 1 plant. The second dataset, which was used to perform large-scale sequence alignment between representative organisms with other species, contained assembled genomes of 16098 bacteria, 11695 viruses and 1496 eukaryotes including 193 mammals, 288 non-mammalian vertebrates, 312 invertebrates, 154 plants, 459 fungi and 90 protozoa. Detailed information about these genomes can be found in Table S12.

### Pipeline to identify HGTs

Figure S1 showed the pipeline to identify HGTs. Firstly, we identified genomic regions distinguishable from the rest of the genome based on k-mer frequencies. The selected genomics regions were then aligned with other genomes. A genomic region was considered as a candidate HGT if its sequence level conservation was discordant to its species phylogenetic tree. Before sequence alignment, we divided all genomes into three different groups based on their taxonomic information, including self group (SG), closely related group (CRG) and distantly related group (DRG). The self group contains organisms whose common ancestor was the one that an HGT event actually occurred. The genomic regions were considered as candidate HGT regions if their sequence conservation levels were discordant to the phylogenetic tree of relevant species. Specifically, genomic fragments were determined as HGT sequence occurring in the SG if they had higher identity percentage with species in DRG than that in CRG. Some similar sequences in DRG of candidate HGT regions were checked by alignment with whole genome sequencing (WGS) raw data of the same organism to avoiding contamination artifacts. We then clustered the HGT sequences to obtain non-redundant HGTs. Phylogenetic trees for related species and homologous sequences were built for each HGT related species and homologous sequences of candidate HGT sequences.

### Sequence composition-based genomic fragments filtering

Due to the large sizes of many reference genomes of model organisms, we first screened the potential genomic regions harboring HGT sequences. For each model organism species, we split the genome sequences into 1000-bp segments with 200-bp overlapped regions across all chromosomes. Sequence segments with Ns were left out. Four-bp kmer frequencies were obtained for the whole genome sequences as well as all genome segments. Euclidean distance was used to measure the difference between each segment and the whole genome sequence. All the distances were sorted in descending order. Finally, the fragments whose distances ranked in the top 10% for human, cow and mouse due to their larger genome sizes, top 20% for the other representative organisms were chosen for further analysis.

In the above statement, two parameters (k-mer size and fragments percentage) needed to be set. We chose appropriate parameters using 299 reported HGTs in the human genome[19], simple repeat sequences of which were filtered out using TRF(version 4.09)[40]. We tried different kmer sizes (1∼6), and k=4 was selected because the highest portion of HGTs previously reported in the human genome were screened out in this situation (Figure S10). A very high portion (>80%) of these human HGTs reported previously were kept in our results even if we only input top 5% of the fragments with highest differences to the human genome (Figure S10).

### Sequence alignment

The sequence alignment was conducted using LASTZ (version 1.04.00)[41] for the filtered fragments of the model organisms and the whole genomes of other species with the following arguments: “--format=axt+ --ambiguous=iupac”.

### Re-screening the fragments and search for HGTs

Every target organism belongs to its own kingdom, phylum, class and order. For each classification level (phylum, class, order and species) of each representative organism, the other organisms were separated into three groups: self group (SG) including all species in the same classification level with target organism, a closely related group (CRG) including all species in the same kingdom with target organism except SG, and a distantly related group (DRG) including all species in the different kingdoms with target organism. For example, when using class as the classification level and using human as the target organism, all mammals were regarded as SG, while the non-mammalian metazoan formed the CRG and DRG including all plants, fungi, protozoa, bacteria and viruses. We further screened the filtered fragments based on alignment results from LASTZ, to identify regions with discordant evolutionary relationships.

The aligned regions (ARs) of the input fragments were retrieved and used to identify putative HGTs. Firstly, we kept ARs that matched to DRG species that were longer than 135bp with a nucleotide identity percentage greater than 50%. For these ARs, we compared the alignment results for CRG species, for which the identity percentage threshold was set to 50%. ARs were regarded as putative HGTs when they had higher similar sequences in DRG species than that in CRG species. To reduce false positive results generated by incorrect alignments, we removed ARs that contained the character ‘N’ or whose GC percentages were less than 0.3 or greater than 0.6. Finally, we checked the repetitive regions overlapping with ARs. RepeatMasker tracks were downloaded from the UCSC Genome Browser, or we ran *de novo* RepeatMasker (version 4.0.7)[42] (http://www.repeatmasker.org) to label the repeats of ARs. We then removed ARs that overlapped with simple repeats or low complexity repeats. We also use TRF(version 4.09)[40] to remove ARs that overlapped with any random repeats. The remaining ARs were considered as candidate HGTs, and were used to build trees to determine discordance with known evolutionary relationships.

### Identifying non-redundant HGTs

HGTs were clustered using the cd-hit-est program (version 4.6.6)[43] with minimum nucleotide identity set at 80%. The longest sequences from each cluster were selected to represent the non-redundant HGTs.

### Validation of similar sequences in other kingdoms with WGS datasets

Discordant HGT trees, constructed from discordant sequences from reference genomes, were the principal evidence for identifying HGTs from our pipeline. Thus, the power for detecting HGTs depended heavily on the quality of the reference genomes. Contaminating sequences from other species were the most likely sources of false positives. For representative organisms, most candidate transferred DNA were also found in their sibling lineages, therefore the probability of sequencing contamination was negligible. However, the inaccurate reference genomes of other organisms (such as parasites and protozoan pathogens) could cause false positive results due to sequencing contamination. For example, if an abnormal HGT tree consists of only one parasite and several primates, and the process of constructing the reference genome of this parasite was contaminated by human DNA, this DNA transfer would be an artifact.

We checked for contamination artifacts in some similar sequences in DRG by alignment with whole genome sequencing (WGS) raw data of the same organism. Firstly, we calculated the distributed of similar sequences of candidate HGTs in other kingdoms. If the number of similar sequences of candidate HGTs in one kingdom is less than 10 (the threshold in bacteria was 50), all similar sequences in this kingdom would be checked by WGS datasets. For each similar sequence, we obtain both upstream and downstream 150bp sequences of its and downloaded multiple WGS raw datasets from the SRA database that were not used to construct the reference genome. Sequence alignment and the calculation of sequencing depth was done using Bowtie2 (version 2.2.4)[44] with default parameters. The sequencing depth of HGTs should not be significantly lower than that of 150bp upstream and downstream sequences of HGTs (using student’s t test, p>0.05) and the sequencing depth of HGTs should be more than 10. Once a sequence got the above standard within at least 2 samples, it was classified as not an artifact. The results are shown in Table S13.

### Exclusion of mitochondrial or chloroplast DNA

Complete mitochondrial genomes of 10 representative organisms and *Arabidopsis thaliana* chloroplast DNA were obtained from NCBI, and we then searched with BLASTN against non-redundant HGTs. With the argument “-evalue 1e-5”, we found no homologous DNA sequences in mitochondrial or chloroplast genomes.

### Remove HGTs present in Endogenous viruses

Endogenous retroviruses (ERVs) are widespread in vertebrates, making up nearly 8% of the genome of *Homo sapiens*[45]. ERVs in human share sequence homology with other primate ERVs[46]. Therefore, in order to avoid reporting sequences as HGTs that are actually from ancestral inheritance, we removed all HGTs found in ERVs. We collected ERVs from the repeat annotation of the UCSC genome browser, except for *Saccharomyces cerevisiae S288C* and *Arabidopsis thaliana*. All HGTs that overlapped with ERV genomic coordinates or aligned to ERVs using BLASTN (identity>90% and length>100bp) were removed.

### Construction of HGT phylogenetic tree

For each HGT, we searched for homologous sequences in other species based on the LASTZ output. When multiple regions in a species met the criteria, the best matched sequence, which had the maximal score weighted by the identity and multiplied by the alignment length, was picked to represent the homologous sequence. Based on the HGT sequence and the homologous sequences collected from other species, we ran multiple sequence alignment using MAFFT (version 7.520)[47] and then trimmed ambiguously aligned regions using trimAl (version 1.4.rev15)[48]. We then used the resulting alignment to infer the ML tree using IQ-TREE (version 2.2.2.3)[49] with its best-fitting model of amino acid evolution and 1000 ultrafast bootstrapping replicates[50]. Finally, we rooted each ML tree at the midpoint using the ape and phangorn R packages[51, 52] and visualized these trees using iTOL (version 6.8.2)[53]. The homologous regions in other species and phylogenetic trees for non-redundant HGTs can be found in Table S14.

### Evaluation of the pipeline using simulated datasets

We constructed a simulated genome (called genome H) with 175 HGTs from a set of distantly related genomes (called Genome set D) to the human genome. Genome set D has 4 cruciferous plant genomes, including *Arabidopsis thaliana*, *Brassica napus*, *Brassica oleracea var. oleracea* and *Brassica rapa*), while Genome set C contains 4 primate genomes, *Pan paniscus*, *Pan troglodytes*, *Pongo abelii* and *Gorilla gorilla gorilla*. The 175 HGTs are sequences that have high similarity with genomes in Genome set D (>90%) but have low similarity (<50%) with genomes in Genome set C, the closely related group of genomes.

Firstly, the genome comparison between genomes in Genome set D was conducted using LASTZ[54] and Multiz[55] to obtain sequences whose identity in all genomes of Genome set D were >90% and lengths >200bps. These sequences were compared with the genomes in Genome set C and the sequences having low similarity (identity <50%) were reserved. The obtained sequences were then clustered using the cd-hit-est program (version 4.6.6)[43] with minimum nucleotide identity set at 80%. The longest sequences from each cluster were selected as simulated HGTs, which were 175 in total. These 175 HGTs were then evenly divided into 10 groups according to their sequence lengths, and the copy numbers of which increased from 2^0^ to 2^9^ (Table S1). Eventually, 175 HGTs with different copy numbers were inserted into the human genome as genome H (Supplementary Data 2). Finally, we ran our pipeline with genome H as the target genome, genome set D as remote genome set, genome set C as closely related genome set and parameters M, N, L as 1, 1, 200 respectively. If the correct HGT region was covered more than 60% of its length by a predicted HGT region, the prediction was considered correct.

### Evaluation of the pipeline using reported whitefly HGTs

We acquired 131 HGTs using our pipeline with *Bemisia tabaci* as target organism and downloaded 170 HGTs previously reported in the whitefly[6]. HGTs identified by our pipeline were considered to be reported if their BLASTN alignment has a matched length>135bp and identity>80%.

### Counting the copy numbers of HGTs

We run BLASTN[56] alignment for non-redundant HGT sequences against their host reference genomes, with the parameter “-e 1e-5”. For each HGT, we selected aligned regions that covered at least 80% of HGT regions with nucleotide identity >80%. We then merged those aligned regions with overlapped coordinates. The copy number of each HGT was determined from the number of merged HGT copies.

### Comparison with reported HGTs in previous studies

We obtained reported HGTs for these model organisms from previous publications, including genomic coordinates and DNA sequences. HGTs in our study were considered novel if they did not match reported HGTs by genomic coordinates or sequence alignment (BLASTN, matched length>135 bp and identity>80%).

### Functional annotation of genes influenced by HGTs

Genome annotation files (GFF or GTF format) were obtained for model organisms from Ensembl [57](http://asia.ensembl.org) and Tair[58] (https://www.arabidopsis.org), and they were used to identify protein-coding genes and non-coding genes likely to be affected by HGTs (overlapping with HGTs with at least 1bp). The Ensembl gene IDs were converted to ENTREZ gene IDs which was input the input of enrichment analysis using DAVID (version 2021)[59] (https://david.ncifcrf.gov). Gene Ontology terms (GO terms) enrichment analysis of these genes were using the clusterProfiler R package [60](FDR<0.05).

## Declarations

### Ethics approval and consent to participate

Not applicable.

### Competing interests

The authors declare that they have no competing interests

### Authors’ contributions

CCW conceived and designed the study. KL, FZY and CCW developed the pipeline and identified HGTs. KL, FZY and ZQD collected the datasets. KL and FZY conducted the visualization. KL, FZY, CCW and DLA wrote the manuscript. KL, FZY, CCW, ZQD and DLA revised the manuscript. All authors read and approved the final manuscript.

## Acknowledgements

This work was supported by grants from Natural Science Foundation of Shanghai (20ZR1428200, 22ZR1433600), National Natural Science Foundation of China (32170643), National key R&D program (2023YFF1001600), Joint International Research Laboratory of Metabolic & Developmental Sciences Joint Research Fund ( MDS-JF-2019A07) and Cross-Institute Research Fund of Shanghai Jiao Tong University (YG2017ZD01). The funders had no role in study design, data collection and analysis, decision to publish, or preparation of the manuscript. T We thank the High Performance Computing Center at Shanghai Jiao Tong University for the computation.

## Data availability

All datasets, supplementary tables and an example of analysis pipeline application are listed in the webpage at http://cgm.sjtu.edu.cn/hgt (password: hgt2019passwd) (this webpage will become freely available after this paper is accepted).

## Code availability

All scripts used in this study are available in GitHub at https://github.com/SJTU-CGM/HGT.git.

### Supplementary Figures, Tables and Datasets

There are 10 Figures, 14 Tables and 2 datasets provided in multiple supplementary files. Descriptions about the figures, tables and datasets are listed below. Supplementary figures are listed in a separate file, while supplementary tables and datasets are accessible from the given URL listed in the data availability.

**Supplementary Figure 1**

The HGT identification system for representative eukaryotes.

**Supplementary Figure 2**

Evaluation of our HGT identification method.

**Supplementary Figure 3**

The distribution of organisms that HGT events occurred.

**Supplementary Figure 4**

Gene ontology (GO) functional enrichment for genes affected by HGTs.

**Supplementary Figure 5**

The distribution of genetic elements affected by HGT regions with different copy numbers in 10 representative organisms.

**Supplementary Figure 6**

Repeat characteristics of genomes of 10 representative organisms.

**Supplementary Figure 7**

HGT examples overlapping with repeat sequences in 6 representative organisms.

**Supplementary Figure 8**

Gene ontology (GO) functional enrichment for protein coding genes affected by HGTs not overlapping with repeat sequences.

**Supplementary Figure 9**

HGT examples overlapping with CDS regions of protein coding genes in 7 representative organisms.

**Supplementary Figure 10**

The impact of parameter setting on the HGT recalling rate for the sequence composition-based filtering step.

**Supplementary Table 1**

Evaluation of the HGT identification pipeline on the simulated dataset.

**Supplementary Table 2**

Evaluation of the HGT identification pipeline on HGTs reported previously.

**Supplementary Table 3**

Detailed information of non-redundant HGTs, including the common ancestors of organisms contained in different self groups (SGs) HGT events occurred, the assembly ID and kingdom of organisms HGT events occurred, copy number in the whole genomes and the number of their overlapping genes.

**Supplementary Table 4**

The distribution of organisms that HGT events occurred.

**Supplementary Table 5**

HGT-appearance numbers between the 10 representative organisms and 1496 eukaryotes.

**Supplementary Table 6**

Gene ontology (GO) functional annotation for genes affected by HGTs.

**Supplementary Table 7**

Repetitive sequence composition of HGTs.

**Supplementary Table 8**

Putative media of horizontal gene transfer of BovB retrotransposons and L1 retrotransposons.

**Supplementary Table 9**

Examples of HGTs in cow, human, Arabidopsis, nematode, yeast and zebrafish genomes.

**Supplementary Table 10**

The influence of HGTs not overlapping with repeat sequence on protein coding genes.

**Supplementary Table 11**

Gene ontology (GO) functional annotation for genes affected by HGTs not overlapping with repeat sequences but overlapping with CDS regions of protein coding genes.

**Supplementary Table 12**

Information of 10 representative organisms and other eukaryotes, bacteria, and viruses.

**Supplementary Table 13**

Coverage matrices of WGS data for HGT homologous sequences in DRG.

**Supplementary Table 14**

Homologous regions in other species and phylogenetic trees for non-redundant HGTs.

**Supplementary Data 1**

Raw output of LASTZ alignment between 10 representative organisms with other eukaryotes (197GB)

URL: http://cgm.sjtu.edu.cn/hgt/data/Supplementary_Data_1.tar

**Supplementary Data 2**

The simulated genome (genome H) with 175 HGTs (2.9GB)

URL: http://cgm.sjtu.edu.cn/hgt/data/Supplementary_Data_2.fa

## Notes

http://cgm.sjtu.edu.cn/hgt

https://github.com/SJTU-CGM/HGT.git

### Competing Interest Statement

The authors have declared no competing interest.

### Summary of Updates

The authors have made extensive revisions, while the main conclusion of this study, that horizontal gene transfer is ubiquitous in eukaryotes, is not changed. 1.Thirteen representative eukaryotes were reduced to 10 representative model organisms with high quality genome sequences. The elephant, lizard and chicken were removed due to their relatively lower genome sequence quality. 2.The genome sequence database used for genome comparison have been updated. 3.All genomes are grouped into three catergories according to their evolutionary distant to the target genome: self group (SG), closely related group (CRG) and distantly related group (DRG). This categorization can make the HGT identification method better understood. 4.We have added systematic evaluation of our HGT identification method, including tests on simulated datasets and comparison with existing method using protein sequence comparison instead of nuclide acid sequence comparison. 5.We compared our HGTs to those newly reported in the past two years and showed that our method is reliable, and HGTs we identified are under more stringent criteria than those used for the reported HGTs.

## References

1. Soucy, S.M., J. Huang, and J.P. Gogarten, Horizontal gene transfer: building the web of life. Nat Rev Genet, 2015. 16(8): p. 472–82.

2. Polz, M.F., E.J. Alm, and W.P. Hanage, Horizontal gene transfer and the evolution of bacterial and archaeal population structure. Trends Genet, 2013. 29(3): p. 170–5.

3. Dagan, T., Y. Artzy-Randrup, and W. Martin, Modular networks and cumulative impact of lateral transfer in prokaryote genome evolution. Proc Natl Acad Sci U S A, 2008. 105(29): p. 10039–44.

4. Cheng, S.F., et al., Genomes of Subaerial Zygnematophyceae Provide Insights into Land Plant Evolution. Cell, 2019. 179(5): p. 1057–1067.

5. Xia, J.X., et al., Whitefly hijacks a plant detoxification gene that neutralizes plant toxins. Cell, 2021. 184(7): p. 1693–1705.

6. Li, Y., et al., HGT is widespread in insects and contributes to male courtship in lepidopterans. Cell, 2022. 185(16): p. 2975-+.

7. Leclercq, S., et al., Birth of a W sex chromosome by horizontal transfer of Wolbachia bacterial symbiont genome. Proceedings of the National Academy of Sciences of the United States of America, 2016. 113(52): p. 15036–15041.

8. Kado, T. and H. Innan, Horizontal Gene Transfer in Five Parasite Plant Species in Orobanchaceae. Genome Biology and Evolution, 2018. 10(12): p. 3196–3210.

9. Lukes, J. and F. Husnik, Microsporidia: A Single Horizontal Gene Transfer Drives a Great Leap Forward. Current Biology, 2018. 28(12): p. R712–R715.

10. Gilbert, C., et al., A role for host-parasite interactions in the horizontal transfer of transposons across phyla. Nature, 2010. 464(7293): p. 1347-U4.

11. Walsh, A.M., et al., Widespread horizontal transfer of retrotransposons. Proceedings of the National Academy of Sciences of the United States of America, 2013. 110(3): p. 1012–1016.

12. Ivancevic, A.M., et al., Horizontal transfer of BovB and L1 retrotransposons in eukaryotes. Genome Biol, 2018. 19(1): p. 85.

13. Huang, J.L., Horizontal gene transfer in eukaryotes: The weak-link model. Bioessays, 2013. 35(10): p. 868–875.

14. Martin, W.F., Too Much Eukaryote LGT. Bioessays, 2017. 39(12).

15. Salzberg, S.L., Horizontal gene transfer is not a hallmark of the human genome. Genome Biol, 2017. 18(1): p. 85.

16. Leger, M.M., et al., Demystifying Eukaryote Lateral Gene Transfer. Bioessays, 2018. 40(5).

17. Keeling, P.J. and J.D. Palmer, Horizontal gene transfer in eukaryotic evolution. Nature Reviews Genetics, 2008. 9(8): p. 605–618.

18. Ivancevic, A.M., et al., Horizontal transfer of BovB and L1 retrotransposons in eukaryotes. Genome Biology, 2018. 19.

19. Huang, W., et al., Widespread of horizontal gene transfer in the human genome. BMC Genomics, 2017. 18(1): p. 274.

20. Sun, B.F., et al., Horizontal functional gene transfer from bacteria to fishes. Scientific Reports, 2015. 5.

21. Pace, J.K., et al., Repeated horizontal transfer of a DNA transposon in mammals and other tetrapods. Proceedings of the National Academy of Sciences of the United States of America, 2008. 105(44): p. 17023–17028.

22. Crisp, A., et al., Expression of multiple horizontally acquired genes is a hallmark of both vertebrate and invertebrate genomes. Genome Biology, 2015. 16.

23. Carr, M., D. Bensasson, and C.M. Bergman, Evolutionary Genomics of Transposable Elements in Saccharomyces cerevisiae. Plos One, 2012. 7(11).

24. Novick, P., et al., Independent and parallel lateral transfer of DNA transposons in tetrapod genomes. Gene, 2010. 449(1-2): p. 85–94.

25. Ma, J.C., et al., Major episodes of horizontal gene transfer drove the evolution of land plants. Molecular Plant, 2022. 15(5): p. 857–871.

26. Jurka, J., et al., Repetitive sequences in complex genomes: Structure and evolution. Annual Review of Genomics and Human Genetics, 2007. 8: p. 241–259.

27. Nowell, F., The blood picture resulting from Nuttallia (=Babesia) rodhaini and Nuttallia (=Babesia) microti infections in rats and mice. Parasitology, 1969. 59(4): p. 991–1004.

28. Doggett, S.L., et al., Bed Bugs: Clinical Relevance and Control Options. Clinical Microbiology Reviews, 2012. 25(1): p. 164-+.

29. Goddard, J. and R. deShazo, Bed Bugs (Cimex lectularius) and Clinical Consequences of Their Bites. Jama-Journal of the American Medical Association, 2009. 301(13): p. 1358–1366.

30. Alexander, W.G., et al., Horizontally acquired genes in early-diverging pathogenic fungi enable the use of host nucleosides and nucleotides. Proc Natl Acad Sci U S A, 2016. 113(15): p. 4116–21.

31. Lukes, J. and F. Husnik, Microsporidia: A Single Horizontal Gene Transfer Drives a Great Leap Forward. Curr Biol, 2018. 28(12): p. R712–R715.

32. Marín, I. and C. Lloréns, retrotransposons:: Description of new elements and evolutionary perspectives derived from comparative genomic data. Molecular Biology and Evolution, 2000. 17(7): p. 1040–1049.

33. Jordan, I.K., L.V. Matyunina, and J.F. McDonald, Evidence for the recent horizontal transfer of long terminal repeat retrotransposon. Proceedings of the National Academy of Sciences of the United States of America, 1999. 96(22): p. 12621–12625.

34. Friesen, N., A. Brandes, and J.S. Heslop-Harrison, Diversity, origin, and distribution of retrotransposons (and) in conifers. Molecular Biology and Evolution, 2001. 18(7): p. 1176–1188.

35. Novikova, O., G. Smyshlyaev, and A. Blinov, Evolutionary genomics revealed interkingdom distribution of Tcn1-like chromodomain-containing Gypsy LTR retrotransposons among fungi and plants. Bmc Genomics, 2010. 11.

36. Rao, W., R.E. Isaac, and J.N. Keen, An analysis of the lipid raft proteome using geLC-MS/MS. Journal of Proteomics, 2011. 74(2): p. 242–253.

37. Ma, X.X., et al., Structures of yeast glutathione--transferase Gtt2 reveal a new catalytic type of GST family. Embo Reports, 2009. 10(12): p. 1320–1326.

38. Langille, M.G.I., W.W.L. Hsiao, and F.S.L. Brinkman, Detecting genomic islands using bioinformatics approaches. Nature Reviews Microbiology, 2010. 8(5): p. 372–382.

39. Pruitt, K.D., T. Tatusova, and D.R. Maglott, NCBI reference sequences (RefSeq): a curated non-redundant sequence database of genomes, transcripts and proteins. Nucleic Acids Research, 2007. 35: p. D61–D65.

40. Benson, G., Tandem repeats finder: a program to analyze DNA sequences. Nucleic Acids Research, 1999. 27(2): p. 573–580.

41. Harris, R.S. Improved pairwise alignment of genomic dna. 2007.

42. Price, A.L., N.C. Jones, and P.A. Pevzner, De novo identification of repeat families in large genomes. Bioinformatics, 2005. 21: p. I351–I358.

43. Li, W. and A. Godzik, Cd-hit: a fast program for clustering and comparing large sets of protein or nucleotide sequences. Bioinformatics, 2006. 22(13): p. 1658–9.

44. Langmead, B. and S.L. Salzberg, Fast gapped-read alignment with Bowtie 2. Nature Methods, 2012. 9(4): p. 357–U54.

45. Paces, J., A. Pavlicek, and V. Paces, HERVd: database of human endogenous retroviruses. Nucleic Acids Research, 2002. 30(1): p. 205–206.

46. Johnson, W.E., Endogenous Retroviruses in the Genomics Era. Annual Review of Virology, Vol 2, 2015. 2: p. 135–159.

47. Katoh, K. and D.M. Standley, MAFFT Multiple Sequence Alignment Software Version 7: Improvements in Performance and Usability. Molecular Biology and Evolution, 2013. 30(4): p. 772–780.

48. Capella-Gutiérrez, S., J.M. Silla-Martínez, and T. Gabaldón, trimAl: a tool for automated alignment trimming in large-scale phylogenetic analyses. Bioinformatics, 2009. 25(15): p. 1972–1973.

49. Nguyen, L.T., et al., IQ-TREE: A Fast and Effective Stochastic Algorithm for Estimating Maximum-Likelihood Phylogenies. Molecular Biology and Evolution, 2015. 32(1): p. 268–274.

50. Minh, B.Q., M.A.T. Nguyen, and A. von Haeseler, Ultrafast Approximation for Phylogenetic Bootstrap. Molecular Biology and Evolution, 2013. 30(5): p. 1188–1195.

51. Paradis, E., J. Claude, and K. Strimmer, APE: Analyses of Phylogenetics and Evolution in R language. Bioinformatics, 2004. 20(2): p. 289–290.

52. Schliep, K.P., phangorn: phylogenetic analysis in R. Bioinformatics, 2011. 27(4): p. 592–593.

53. Letunic, I. and P. Bork, Interactive tree of life (iTOL) v3: an online tool for the display and annotation of phylogenetic and other trees. Nucleic Acids Research, 2016. 44(W1): p. W242–W245.

54. Harris, R.S., Improved pairwise alignment of genomic dna. 2007, The Pennsylvania State University.

55. Blanchette, M., et al., Aligning multiple genomic sequences with the threaded blockset aligner. Genome Res, 2004. 14(4): p. 708–15.

56. Camacho, C., et al., BLAST plus : architecture and applications. Bmc Bioinformatics, 2009. 10.

57. Zerbino, D.R., et al., Ensembl 2018. Nucleic Acids Research, 2018. 46(D1): p. D754–D761.

58. Poole, R.L., The TAIR database. Methods Mol Biol, 2007. 406: p. 179–212.

59. Huang, D.W., B.T. Sherman, and R.A. Lempicki, Systematic and integrative analysis of large gene lists using DAVID bioinformatics resources. Nature Protocols, 2009. 4(1): p. 44–57.

60. Yu, G.C., et al., clusterProfiler: an R Package for Comparing Biological Themes Among Gene Clusters. Omics-a Journal of Integrative Biology, 2012. 16(5): p. 284–287.

